# Lysosomal perturbations in dopaminergic neurons derived from induced pluripotent stem cells with *PARK2* mutation

**DOI:** 10.1101/734244

**Authors:** Justyna Okarmus, Helle Bogetofte, Sissel Ida Schmidt, Matias Ryding, Silvia Garcia Lopez, Alberto Martínez-Serrano, Poul Hyttel, Morten Meyer

**Author notes:** Authors contributed equally to this work (Justyna Okarmus, Helle Bogetofte). Corresponding author: Morten Meyer, Ph.D., Department of Neurobiology Research, Institute of Molecular Medicine, University of Southern Denmark, J.B. Winsløws Vej 21, st, DK-5000 Odense C, Denmark.

## Abstract

Mutations in the *PARK2* gene encoding parkin, an E3 ubiquitin ligase, are associated with autosomal recessive early-onset Parkinson’s disease (PD). While parkin has been implicated in the regulation of mitophagy and proteasomal degradation, the precise mechanism leading to neurodegeneration in both sporadic and familial PD upon parkin loss-of-function mutations remains unknown. Cultures of isogenic induced pluripotent stem cell (iPSC) lines with and without *PARK2* knockout (KO) enable mechanistic studies of the effect of parkin deficiency in human dopaminergic neurons. In the present study, we used such cells to investigate the impact of *PARK2* KO on the lysosomal compartment combining different approaches, such as mass spectrometry-based proteomics, electron microscopy (TEM) analysis and functional assays. We discovered a clear link between parkin deficiency and lysosomal alterations. *PARK2* KO neurons exhibited a perturbed lysosomal morphology, displaying significantly enlarged and electron-lucent lysosomes as well as an increased total lysosomal content, which was exacerbated by mitochondrial stress. In addition, we found perturbed autophagic flux and decreased lysosomal enzyme activity suggesting an impairment of the autophagy-lysosomal pathway in parkin-deficient cells. Interestingly, activity of the GBA-encoded enzyme, β-glucocerebrosidase, was significantly increased suggesting the existence of a compensatory mechanism. In conclusion, our data provide a unique characterization of the morphology, content, and function of lysosomes in *PARK2* KO neurons, thus revealing a new important connection between mitochondrial dysfunction and lysosomal dysregulation in PD pathogenesis.

## Introduction

Parkinson’s disease (PD) is a progressive neurodegenerative disorder affecting 1-2% of the population. Although the majority of PD patients develop late-onset sporadic disease, a subpopulation of patients develops early-onset or familial PD forms associated with various genetic mutations. Studies of the effects of these mutations can provide valuable insights into the molecular pathways and mechanisms that lead to degeneration of dopaminergic neurons in PD (1–4).

Mutations in the *PARK2* gene, encoding the protein parkin, have been identified as the most common cause of autosomal recessive early-onset PD and have underlined the importance of mitochondrial dysfunction in PD pathogenesis (5–7). Parkin is a multifunctional E3 ubiquitin ligase involved in several cellular processes. Parkin-mediated ubiquitination of mitochondrial proteins (8–11) triggers clearance of impaired mitochondria through the autophagy-lysosome pathway (ALP) (12). Lysosomes are organelles specialized for the degradation of macromolecules derived from the extracellular space through endocytosis or phagocytosis, or from the cytoplasm through autophagy. In recent years, the importance of lysosomes in pathology has been indicated by a rapidly growing number of human disorders linked to defects in lysosomal function including PD (13, 14) in which the accumulation of non-degraded and misfolded proteins occurs. Mutations in the GBA gene, coding for the lysosomal glycohydrolase β-glucocerebrosidase (GCase), cause Gaucher’s disease and several studies have reported GBA mutations as the numerically greatest genetic risk factor for PD (15–17). A number of studies point to an interplay between mitochondrial homeostasis and proper lysosomal function. Diseases caused by mutations of ALP proteins often exhibit mitochondrial defects as well (18–20). Of relevance for PD, loss of GCase activity leads to mitochondrial dysfunction indicating that impaired lysosomal function negatively impacts mitochondria (17). Supporting this, autophagy enhancing drugs such as rapamycin has neuroprotective effects against the mitochondrial complex I inhibitor rotenone in cellular models of PD (21). Interestingly, mitochondrial dysfunction induced by rotenone treatment alters the expression of lysosomal genes, which perhaps is explained by the fact that mitophagy induction through nuclear translocation of transcription factors regulates mitochondrial and lysosomal biogenesis (22, 23). Recent studies have documented mitochondria-lysosome membrane contact sites, which enable bidirectional regulation of mitochondrial and lysosomal dynamics, and have demonstrated how mitochondrial impairment supresses autophagic flux, overall suggesting a complex mutual relationship between these two cellular compartments (24–28).

However, the exact relationship between mitochondrial- and lysosomal function in PD is not well defined (23, 27, 28) and elucidation of a potential link between them and its role in the pathogenic process is still required. By studying the lysosomal compartment and function in the context of parkin deficiency, we sought to address whether chronic mitochondrial dysfunction causes lysosomal impairment, contributing to PD pathogenesis. For this purpose, we studied isogenic iPSC-derived neuronal cultures with and without *PARK2* mutation, which we have recently demonstrated lead to several mitochondrial defects (29). Parkin deficiency resulted in a number of perturbations including altered lysosomal content, morphology and function as well as impaired autophagic flux, indicating a connection between parkin deficiency and lysosomal disturbances.

## Results

### Identical differentiation potentials of *PARK2* KO iPSCs and control lines

To study the disease mechanism underlining *PARK2*-mediated PD, we analyzed two isogenic iPSC lines created from a healthy control iPSC line, where KO of the *PARK2* gene was created by zinc finger nuclease gene editing technology (30). Detailed cell line information is reported in our recent study (29).

*PARK2* KO and isogenic control iPSC-derived neuronal stem cells (NSCs) were differentiated simultaneously to assess the efficiency of midbrain dopaminergic neuron yield (*Fig. 1A*). *Fig. 1B* shows representative immunofluorescence pictures of cultures differentiated for 25 days, revealing a large percentage of MAP2+ mature neurons with distinct cell bodies and long branched processes forming highly interconnected networks. No apparent difference in the percentage of mature neurons was observed (control: 69.8 ± 1.0%, *PARK2*: 68.4 ± 0.9%) (*Fig. 1C*). The differentiated cells were also positive for NeuN and synaptophysin markers, which confirmed their maturity (<u>Bogetofte et al., submitted</u>). Many of the generated neurons were found to co-localize with the dopaminergic marker tyrosine hydroxylase (TH), which is the rate-limiting enzyme in the production of dopamine. No apparent difference in the amount of TH+ dopaminergic neurons was observed between cell lines (control: 25.25 ± 1.0%, *PARK2*: 24.6 ± 1.1%) (*Fig. 1D*). Moreover, the presence of GABAergic+ neurons and a small population of GFAP+ astrocytes was also found in the cultures (<u>Bogetofte et al., submitted</u>). Western blot analysis showed similar amounts of MAP2 and TH protein expression between control and *PARK2* KO cell lines, confirming the immunofluorescence staining (*Figs. 1E, F*). qRT-PCR analysis detected the presence of midbrain dopaminergic specific markers (EN1, NURR1, GIRK2) in the differentiated neuronal cultures (*Fig. 1G*). Both lines showed comparable expression levels with no significant differences. These data show that *PARK2* KO does not affect the neuronal differentiation potential of the iPSC-derived NSCs, as both the *PARK2* KO and isogenic control lines were equally efficient in generating midbrain dopaminergic neurons.

**Figure 1:**
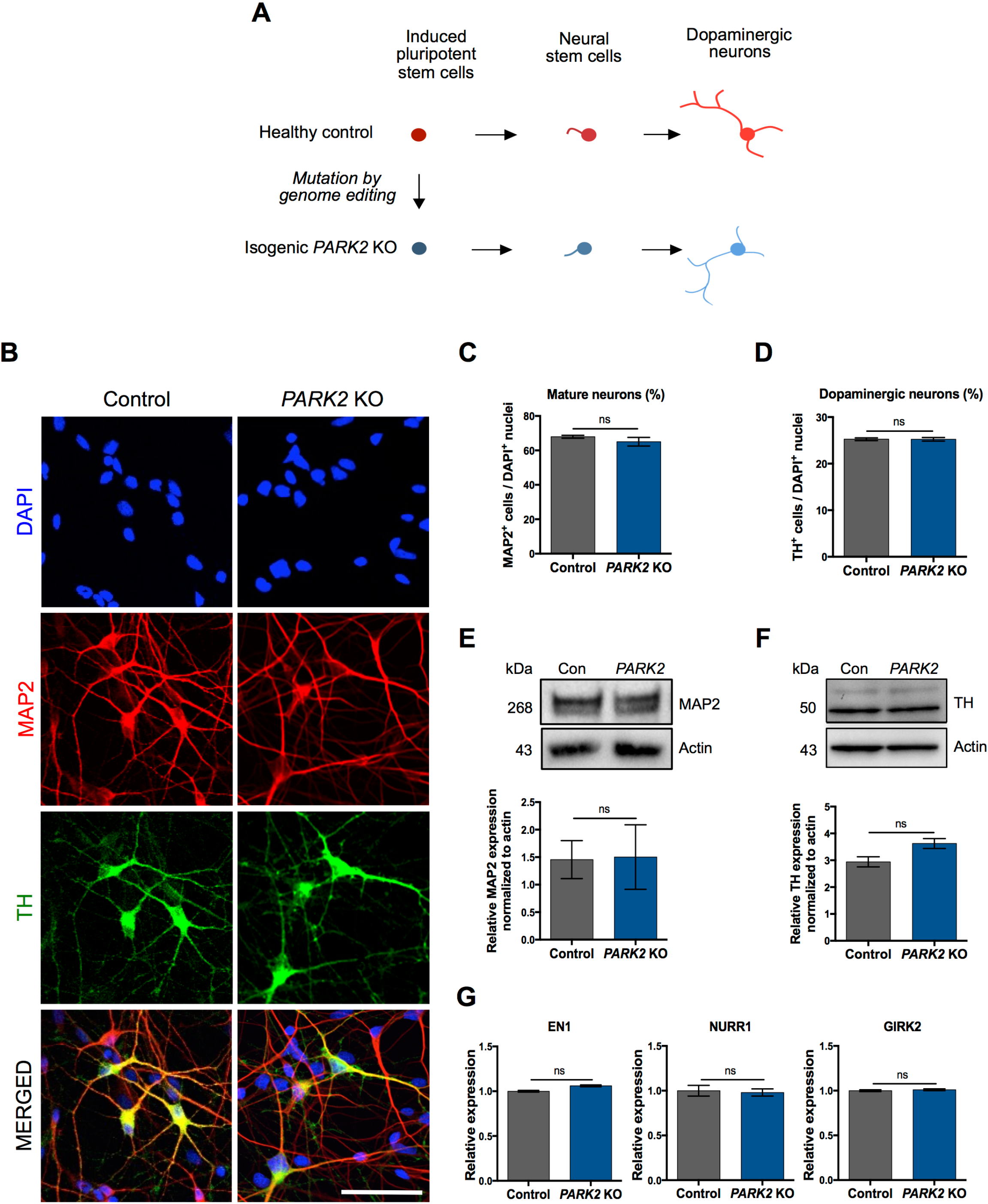
General characterization of neurons derived from *PARK2* KO and healthy isogenic induced pluripotent stem cells (iPSCs). A) Graphical overview of the differentiation of iPSCs to neural stem cells (NSCs) and fully committed dopaminergic neurons. B) Immunofluorescence analysis of MAP2 (mature neurons, red), and TH (dopaminergic neurons, green) expression in *PARK2* KO iPSC and control iPSC lines at day 25 of differentiation. Cell nuclei are visualized using DAPI (blue). Representative pictures are shown. Scale bar: 50 μm. C-D) Quantitative assessment of C) MAP2+ mature neurons and D) TH+ dopaminergic neurons in *PARK2* KO and control iPSC lines. All data are presented as mean ± SEM, n=15; 5 independent differentiations. E-F) Western blotting and densitometry analysis of E) MAP2 and F) TH protein expression levels in *PARK2* KO and control iPSC lines. Protein expression levels were normalized to α-actin. Data are presented as mean ± SEM; 3 independent experiments. G) qPCR analysis for midbrain/dopaminergic markers (EN1, NURR1 and GIRK2). GAPDH, 18S, HPRT were used as endogenous references. Data were normalized to control levels and presented as mean ± SEM, 2 independent experiments.

### Proteomic changes and increased lysosomal content in *PARK2* KO neurons

As earlier reported we have subjected *PARK2* KO and isogenic control neuronal cultures to a mass spectrometry-based proteomic analysis, which enabled the identification and quantification of a large number of proteins (29). Based on this recently published dataset (29) we detected significant changes in levels of 22 lysosomal proteins in *PARK2* KO neurons (*Table 1*). A number of these proteins were of importance for vesicle-mediated protein trafficking through the endosomal-lysosomal system and for the ALP pathway (*Table 1*). This indicated that lysosomal perturbations were indeed present in the *PARK2* KO neurons and led us to examine the overall lysosomal content of *PARK2* KO neurons. Interestingly, levels of both lysosomal associated membrane protein 1 and 2a (LAMP1/2a), which are often used as general markers for lysosomes, were significantly elevated as quantified by Western blotting (*Figs. 2A, B)*, pointing to an overall increased lysosomal content in the *PARK2* KO neurons.

**Table 1:** Changes in lysosomal content in *PARK2* KO neurons revealed by proteomic analysis. Table listing lysosomal proteins identified by proteomic analysis with the ratio of their protein levels in *PARK2* KO neurons compared to controls; q-value (FDR-adjusted p-value) and number of unique peptides, n = 3, three independent differentiations.

| Accession | Gene name | Description | *PARK2* KO / Control | q-value | Unique Peptides |
| --- | --- | --- | --- | --- | --- |
| P27449 | ATP6V0C | V-type proton ATPase 16 kDa proteolipid subunit | **1.56** | 0.005 | 4 |
| Q6IQ22 | RAB12 | Ras-related protein Rab-12 | **1.31** | 0.014 | 9 |
| Q9Y5W7 | SNX14 | Sorting nexin-14 | **1.30** | 0.018 | 2 |
| Q8IY95 | TMEM192 | Transmembrane protein 192 | **1.25** | 0.025 | 2 |
| O15400 | STX7 | Syntaxin-7 | **1.23** | 0.036 | 13 |
| Q96GC9 | VMP1 | Vacuole membrane protein 1 | **1.21** | 0.035 | 3 |
| Q9H267 | VPS33B | Vacuolar protein sorting-associated protein 33B | **0.87** | 0.039 | 7 |
| P78537 | BLOC1S1 | Biogenesis of lysosome-related organelles complex 1 subunit 1 | **0.85** | 0.042 | 3 |
| Q969J3 | BORCS5 | BLOC-1 related complex subunit 5 | **0.85** | 0.037 | 2 |
| Q9H305 | CDIP1 | Cell death-inducing p53-target protein 1 | **0.84** | 0.028 | 2 |
| P07858 | CTSB | Cathepsin B | **0.84** | 0.047 | 2 |
| Q9UN37 | VPS4A | Vacuolar protein sorting-associated protein 4A | **0.83** | 0.036 | 8 |
| P86790 | CCZ1B | Vacuolar fusion protein CCZ1 homolog B | **0.82** | 0.020 | 3 |
| Q8TAF3 | WDR48 | WD repeat-containing protein 48 | **0.81** | 0.042 | 8 |
| O14964 | HGS | Hepatocyte growth factor-regulated tyrosine kinase substrate | **0.79** | 0.015 | 5 |
| P61073 | CXCR4 | C-X-C chemokine receptor type 4 | **0.79** | 0.016 | 3 |
| O43237 | DYNC1LI2 | Cytoplasmic dynein 1 light intermediate chain 2 | **0.74** | 0.017 | 19 |
| Q01484 | ANK2 | Ankyrin-2 | **0.72** | 0.009 | 167 |
| P61916 | NPC2 | NPC intracellular cholesterol transporter 2 | **0.69** | 0.020 | 3 |
| P60520 | GABARAPL2 | Gamma-aminobutyric acid receptor-associated protein-like 2 | **0.67** | 0.004 | 6 |
| P98164 | LRP2 | Low-density lipoprotein receptor-related protein 2 | **0.63** | 0.003 | 16 |
| Q9H492 | MAP1LC3A | Microtubule-associated proteins 1A/1B light chain 3A | **0.57** | 0.003 | 2 |

**Figure 2:**
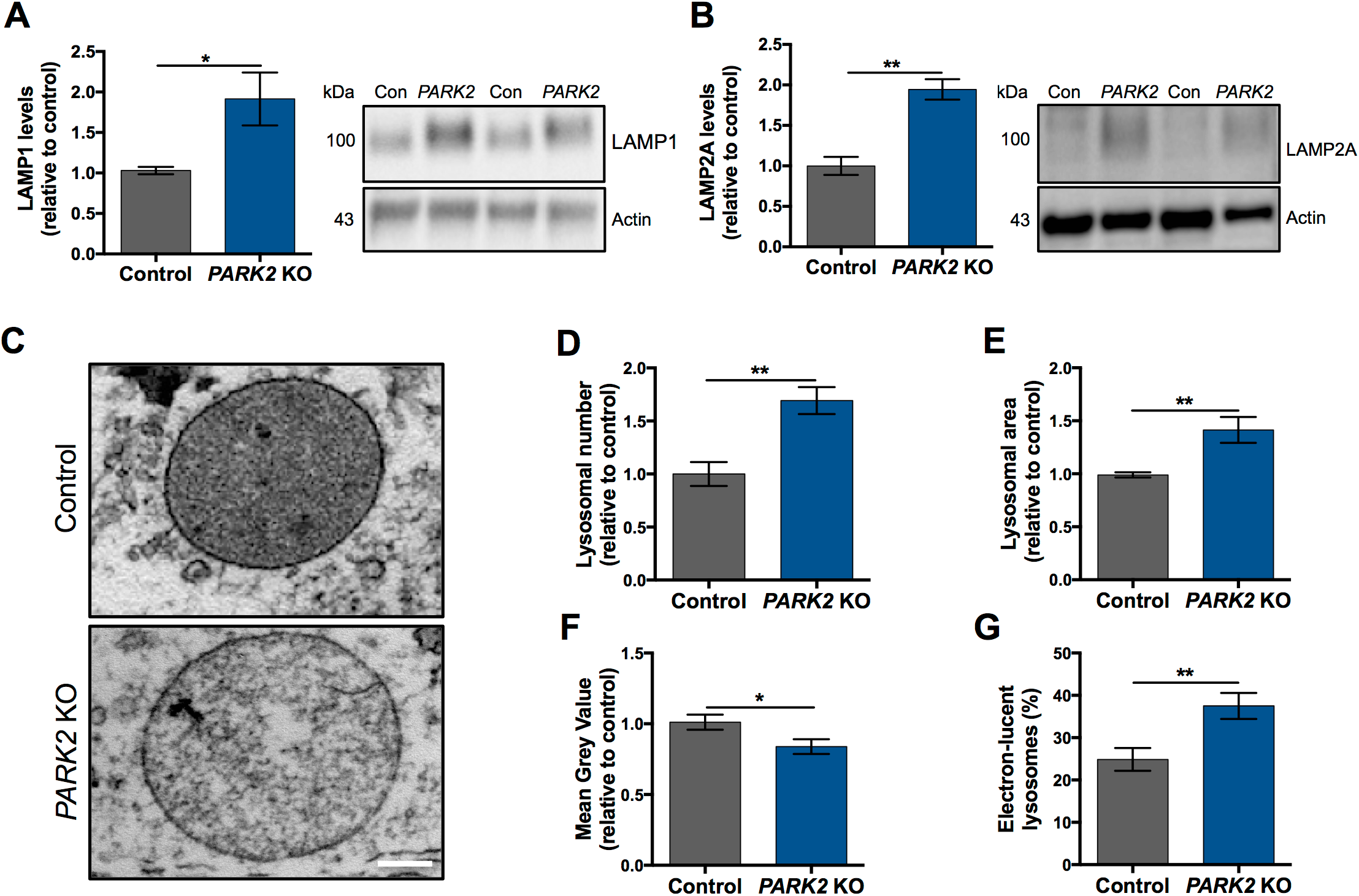
*PARK2* KO neurons exhibit aberrant lysosomes. A, B) Western blotting analysis of the abundance of lysosomal markers B) LAMP1 and C) LAMP2a. Expression levels were normalized to β-actin and are shown relative to control neurons. Data are presented as mean ± SEM of at least 3 independent differentiations. *p < 0.05, and **p < 0.01, analyzed using Student’s t-test. C) Representative TEM micrographs showing the ultrastructure of lysosomes in healthy control (upper image) and *PARK2* KO (bottom image) neurons. Scale bar: 100 nm. D) Quantification of the lysosomal number per TEM micrograph showing lysosomal accumulation in *PARK2* KO neurons. E) Lysosomal size, as a measurement of average lysosomal area, was increased in *PARK2* KO neurons. F) Mean Grey Value of individual lysosomes was reduced in *PARK2* KO neurons. G) Percentage of electron-lucent lysosomes among all lysosomes in a given neuronal population. Data are presented as mean ± SEM, n=6 TEM grids, 3 independent differentiations (in total 45 TEM micrograps per cell line were analyzed. *p < 0.05, and **p < 0.01, analyzed using Student’s t-test.

### TEM evidences for lysosomal abnormalities in *PARK2* KO neurons

To further investigate the lysosomal perturbations suggested by the proteomic analysis we applied transmission electron microscopy (TEM) analysis, which revealed several ultrastructural disturbances within lysosome-like structures. Lysosomes were identified as electron-dense small spherical organelles, enclosed by a single membrane, with a size of about 0.5-1.0 μm in diameter (31). Ultrastructural assessment of the *PARK2* KO and healthy control neurons detected clear differences in lysosomal morphology between the two cell lines. Lysosomes in neurons with *PARK2* KO were more heterogeneous and characterized by the presence of large translucent areas, whereas lysosomes from healthy control neurons more uniformly displayed an electron-dense inner compartment (*Fig. 2C*). Quantitative analysis of the number of lysosomes confirmed a significant accumulation of lysosomes in *PARK2* KO neurons compared to healthy isogenic controls (70% increase, p < 0.01) (*Fig. 2D*). The area of the individual lysosomes was increased for *PARK2* KO neurons by approximately 40% (p < 0.01) compared to controls (*Fig. 2E*). This verified the presence of enlarged lysosomal vacuoles observed for *PARK2* KO neurons. In addition, the mean gray value (a measure of lysosomal electron density) was significantly reduced for *PARK2* KO neurons (20% decrease, p < 0.05) (*Fig. 2F*). This led us to categorize lysosomes into two groups: electron-lucent (lighter) and electron-dense (darker), and we found a significantly greater amount of electron-lucent lysosomes in the *PARK2* KO neurons compared to controls (37.5% vs. 24.8%, respectively, p < 0.01) (*Fig. 2G*). Overall, TEM analysis revealed significant morphological changes in lysosomes between *PARK2* KO neurons and isogenic control neurons. Lysosomal vacuoles appeared to be more abundant, larger and more electron-lucent (lighter) in *PARK2* KO neurons than in controls.

### Altered lysosomal abundance in mature *PARK2* KO neurons

In order to investigate the time course of the lysosomal accumulation during the differentiation of *PARK2* KO neurons, we investigated the abundance of lysosomes at different time points by immunofluorescence staining using the lysosomal marker LAMP1 (*Figs. 3A-E*). We did not observe significant differences in the pattern of LAMP1 staining between *PARK2* KO and isogenic control neurons at the early stages of differentiation day 0 (neural stem cell stage) and 10 (progenitor cell stage) (*Figs. 3A, B, F*). After 25 days of differentiation, when approximately 70% of the neurons were mature (*Fig. 1C*), *PARK2* KO neurons contained significantly increased area of lysosomes per cell, consistent with the obtained TEM data. This was evident from an approximately 25% increase in the area of LAMP1+ staining per DAPI+ nuclei in *PARK2* KO neurons compared to control neurons (*Figs. 3C, F*). The lysosomal area was increasing for both cell lines proportional to the time of differentiation, retaining the observed lysosomal accumulation in *PARK2* KO neurons (*Figs. 3A-F*). Moreover, at the later stages of differentiation in healthy control neurons, lysosomes were well resolved as puncta (*Figs. 3C-E, upper panel, white arrows*). In contrast, lysosomes appeared enlarged and clustered in *PARK2* KO cells (*Figs. 3C-E*, *lower panel, red arrows*). Altogether, our findings demonstrate that parkin dysfunction causes lysosomal accumulation and altered subcellular distribution in mature human *PARK2* KO neurons in a time dependent manner.

**Figure 3:**
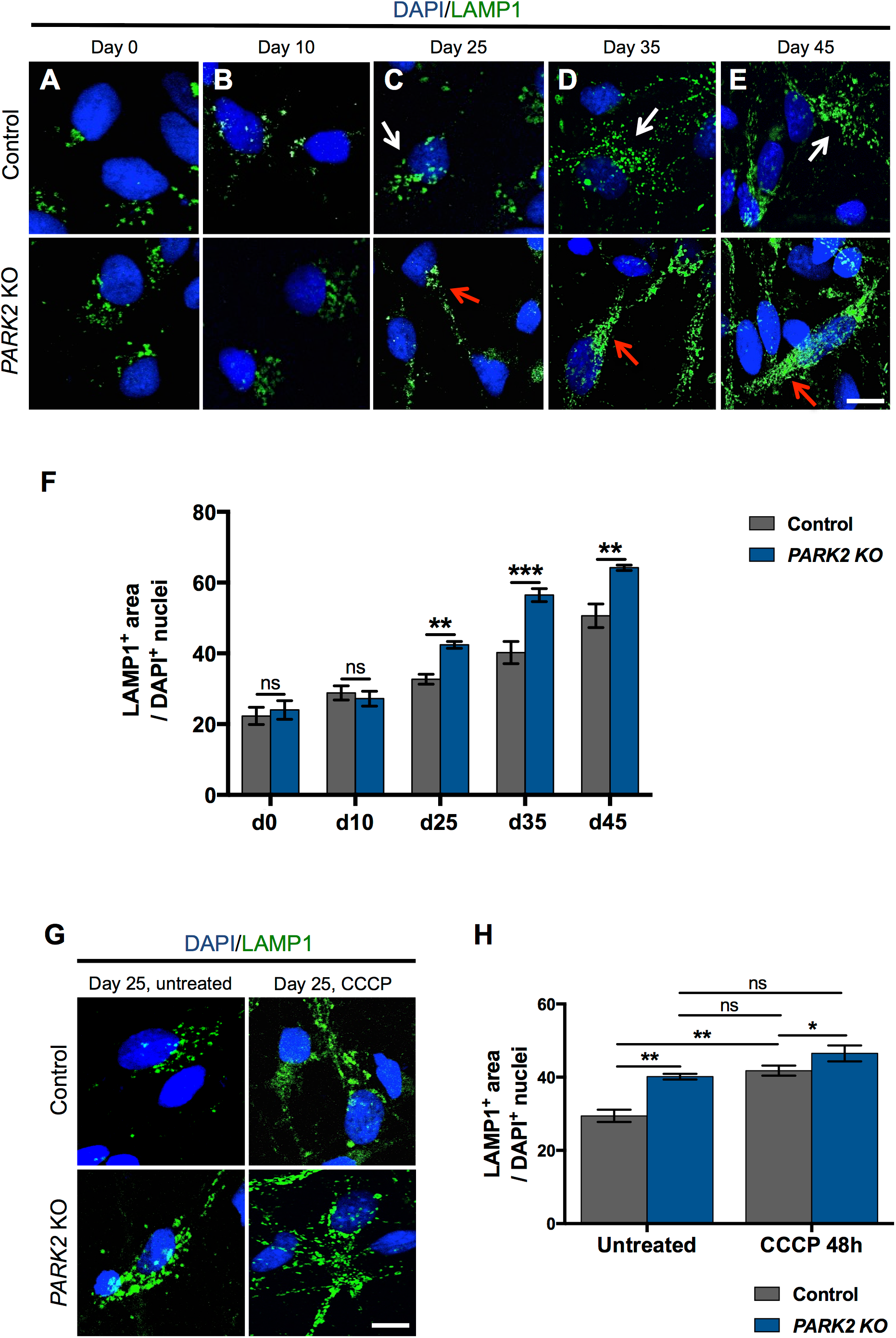
Increased lysosomal area caused by parkin deficiency and CCCP treatment. A-E) Temporal changes in LAMP1 (green) immunoreactivity in iPSC-derived neurons from a healthy control and *PARK*2 KO cells. Cell nuclei were visualized using DAPI (blue). Scale bar: 20 *μ*m. F) Quantification of the area of LAMP1+ lysosomes normalized to the number of DAPI+ nuclei showing no difference between *PARK2* KO and control cells at days 0 and 10, while a significant increase was observed at day 25 and later time points. Data are presented as mean ± SEM of 3 independent differentiations. **p < 0.01, ***p < 0.001, ns: not significant, analyzed with Student’s t-test. G) LAMP1 (green) and DAPI (blue) immunofluorescence staining of *PARK2* KO neurons and isogenic controls at day 25 after CCCP treatment (10 *μ*M, 48h). Scale bar: 20 *μ*m. H) Significant increase of lysosomal area in both control and the *PARK2* KO neurons after CCCP exposure. Data presented as mean ± SEM, three independent differentiations. *p < 0.05, **p < 0.01, ns: not significant, analyzed using one-way ANOVA.

### Increased lysosomal abundance after CCCP treatment

To investigate whether the accumulation of lysosomes observed in *PARK2* KO neurons at day 25 was a direct consequence of mitochondrial impairment we studied the lysosomal response to mitochondrial stress caused by *carbonyl cyanide m-chlorophenyl hydrazine* (CCCP), a proton ionophore that collapses the proton electrochemical gradient across membranes, thus acting as a potent uncoupler of respiratory chain function and oxidative phosphorylation (32). After 48 hours of CCCP (10 μM) exposure, the area of lysosomes in control cells had increased to a level similar to that of *PARK2* KO untreated neurons (41.8 vs. 40.15 arbitrary units, respectively, p = 0.98), strongly indicating that mitochondrial function impairment leads to lysosomal accumulation. Interestingly, CCCP treatment of *PARK2* KO neurons resulted in further enhancement of LAMP1 staining (*Figs. 3G, H*), consistent with the hypothesized mitochondrial dysfunction present in these cells and indicating that this is a general consequence of mitochondrial dysfunction. Taken together, our data show that exposure to mitochondrial stressors, like CCCP, results in elevated lysosomal area in both *PARK2* KO and healthy neurons, which provides further evidence of a causative link between mitochondrial dysfunction and lysosomal alterations.

### Perturbed lysosomal function and autophagic flux in *PARK2* KO neurons

To determine whether the alterations in lysosome protein levels and overall organelle structure may impact lysosomal function, we performed several functional assays. The first analysis assessed lysosomal intracellular activity (general enzyme activity) and revealed a significant > 30% reduction (p < 0.05) in overall functional activity of lysosomes in *PARK2* KO neurons compared to controls (*Figs. 4A, B*). To further investigate the impairment of lysosomal function, we additionally monitored the activity of two essential lysosomal enzymes, β-galactosidase (β-Gal, *Figs. 4C, D*) and β-glucocerebrosidase (GCase, *Fig. 4E-G*). β-Gal activity was linear over the assay time and a clear reduction was observed for the *PARK2* KO neurons when compared to controls, which pattern was more similar to the positive control included in the assay (*Fig. 4C*). Moreover, the specific β-Gal activity was calculated, which confirmed a reduction of activity in the *PARK2* KO neurons (control: 56 ± 5.7 enzymatic units, *PARK2*: 37.5 ± 2.3 enzymatic units, p < 0.01) (*Fig. 4D*). Interestingly and in parallel, the activity of the lysosomal enzyme GCase, encoded by the PD-related *GBA* gene, was significantly increased in the *PARK2* KO neurons (approximately 25% increase, p < 0.001) (*Fig. 4E*). As the protein levels of GCase were not significantly increased (*Fig. 4F*), the enhanced GCase activity was not simply an effect of increased lysosomal numbers or enlargement. Rather it may reflect a compensatory functional activation (*Figs. 4E-G*). In order to establish if the alterations detected in the lysosomal function also affected the autophagic process, as alluded to by the proteomic analysis (*Table 1*), we analyzed through Western blotting the expression of microtubule-associated proteins 1A/1B light chain 3 (gene MAP1LC3A in *Table 1*), a known marker of autophagy (33). The total amount of LC3 protein was significantly decreased in *PARK2* KO neurons at baseline confirming the results from the proteomic analysis *(Table 1*, *Figs. 4H, 4I)*. LC3-II protein turnover was measured as the ratio between protein levels of LC3-II and LC3-I in the presence and absence of bafilomycin A1, which inhibits lysosomal acidification and fusion of autophagosomes with the lysosome. At baseline the *PARK2* KO neurons compared to control had a significantly increased LC3-II/LC3-I ratio, which however was unaffected by bafilomycin A1 treatment. In contrast the control neurons showed a significant increase in the LC3-II/LC3-I ratio upon bafilomycin A1 treatment *(Figs. 4H, 4J)*. This indicates that autophagy induction is occurring, but that in the *PARK2* KO neurons the autophagic process is stalled due to lysosomal impairment. Overall, the *PARK2* KO neurons showed a significantly decreased autophagy process compared to healthy control neurons (approximately 30% decrease, p < 0.01) (*Fig. 4K*). In conclusion, these data overall provide strong and consistent evidence of a perturbed lysosomal activity and compromised autophagic flux in the *PARK2* KO neurons.

**Figure 4:**
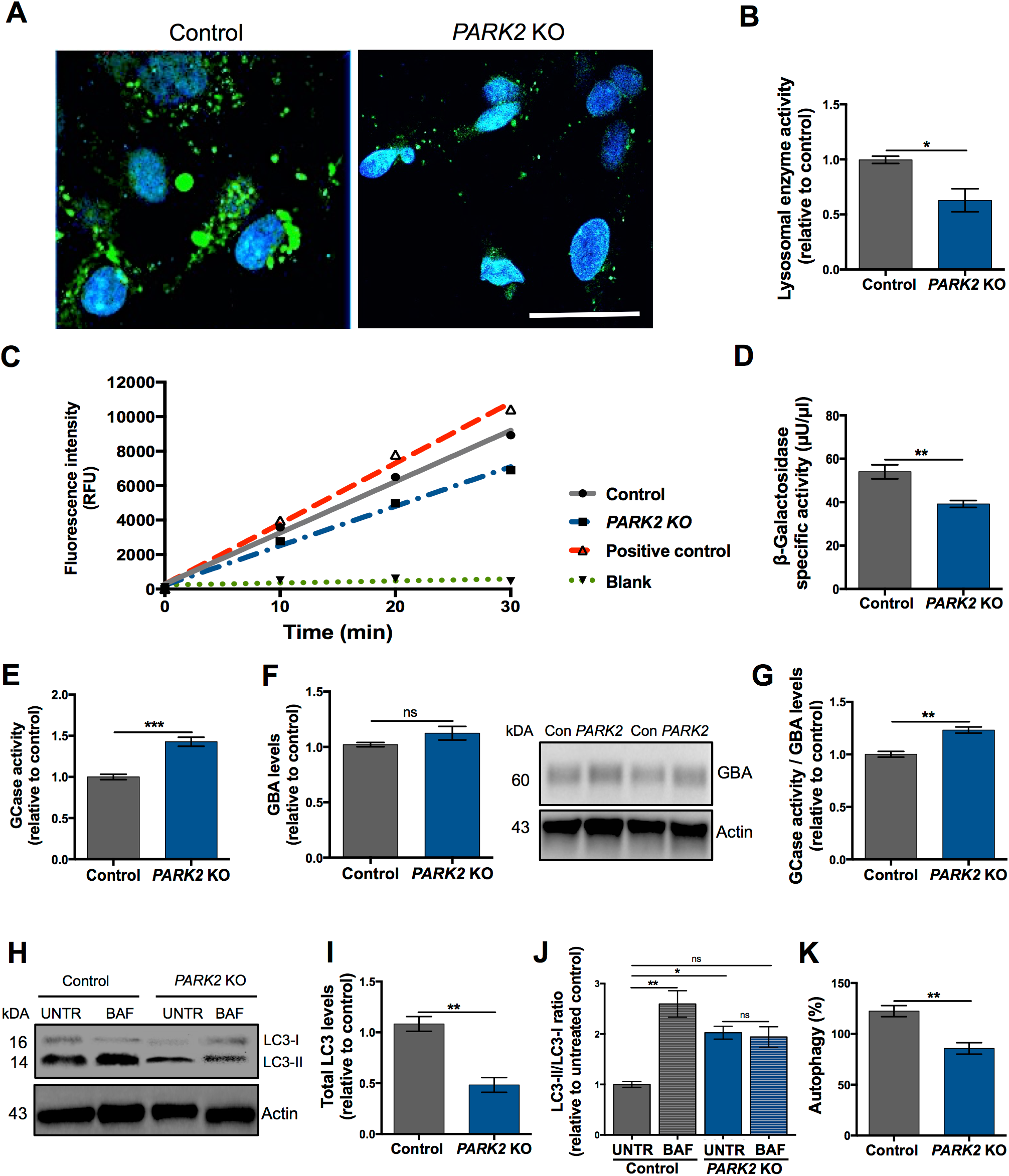
Perturbed lysosomal function and altered autophagic flux in *PARK2* KO neurons. A) General intracellular lysosomal enzyme activity manifested by generation of the fluorescence signal in *PARK2* KO and control neurons. Scale bar: 50 μm. B) Quantification of the signal intensity showed a significant decrease in the overall functional activity of lysosomal enzymes in the *PARK2* KO neurons compared to controls. C) Kinetics of enzymatic activity of β-galactosidase (β-Gal) based on relative fluorescence intensity versus time. Graph represents changes in the *PARK2* KO neurons (blue dashed line) and control neurons (grey solid line), positive control (red dashed line), and blank (green dot-dash line). D) The specific β-Gal activity was significantly reduced for the *PARK2* KO neurons. E) Glucocerebrosidase (GCase) enzyme activity was significantly increased for *PARK2* KO neurons compared to controls. F) Western blotting showed no changes in GCase levels. Expression levels were normalized to β-actin and shown relative to control neurons. G) GCase enzyme activity normalized to GCase levels was significantly increased in the *PARK2* KO neurons. (H-K) Alteration of autophagy in *PARK2* KO neurons. H) Differentiated cells were incubated with 10 nM bafilomycin A1 (BAF) for 6h or with equivalent amount of DMSO as vehicle (UNTR) and western blotting was performed for LC3. I) Total LC3 (LC3-I + LC3-II) levels and J) LC3-II/LC3-I ratios for each group was calculated by densitometric analysis and normalized to the level of α-actin. K) The autophagic flux manifested as the ratio of LC3 between BAF and UNTR of the same group. All data presented as mean ± SEM, 2-3 independent differentiations. *p < 0.05, **p< 0.01, ***p < 0.001, ns: not significant, analyzed using Student’s t-test.

## Discussion

The importance of parkin dysfunction in familial and sporadic PD is well established, but the exact mechanism and pathways are not well understood (6, 33). Parkin was associated with mitochondrial function for the first time through studies in *Drosophila M.* (34). However, today we know that the parkin protein has multiple roles and that it is crucial for the proper function of not only mitochondria but also lysosomes (23, 34). A common feature shared by these two organelles is that mitochondrial defects and lysosomal function impairment can lead to neurological pathologies, suggesting functional connections between these two organelles (35, 36).

In our recent report we demonstrated prominent mitochondrial-related defects and the mitochondrial proteome perturbations (29) in response to parkin deficiency. Therefore, to explore the reciprocal relationship between mitochondrial and lysosomal pathways in PD, in the present study we investigated the impact of parkin dysfunction on lysosome structure and function by applying human iPSC-derived neurons with *PARK2* mutations and isogenic healthy control. We identified striking differences in lysosomal abundance, morphology, content and activity in the *PARK2* KO neurons. Endo-lysosomal alterations have been documented in other parkin dysfunction models (37, 38). However, to our knowledge, this is the first experimental evidence obtained from human *PARK2* KO iPSC-derived neurons.

One of the consequences of mitochondrial deficits caused by parkin dysfunction in this cellular model of PD was an increased lysosomal number and size. This altered phenotype appeared at differentiation day 25, when the cells also exhibit aberrant mitochondria, indicating increased vulnerability of mature neurons (29). Similar lysosomal alterations to those observed in our study were detected in mitochondrial mutants exhibiting disrupted mitochondrial fission-fusion processes and cristae structure and mitochondrial impairment (27, 39–41). A recent study in mice and cellular models of mitochondrial dysfunction reported a ROS-dependent increase in numbers of lysosomal vacuoles, suggesting a possible mechanism (27). Mitochondrial activity is required to maintain proper lysosomal structure and function, as demonstrated by studies applying chemical inhibition of the electron transport chain, as well as in *in vitro* and *in vivo* genetic models of mitochondrial dysfunction. Specifically, disruption of mitochondrial function causes the accumulation of enlarged endo-lysosomal structures (27, 39, 42, 43).

Our study supports these observations, as lysosomal numbers were increased following uncoupling of mitochondrial function by CCCP treatment in both neuronal populations. CCCP-treated control neurons contained almost equal numbers of lysosomes as *PARK2* KO untreated neurons, pointing to an association between lysosomal accumulation and mitochondrial membrane depolarization caused by CCCP. In line with this, an increase in lysosomal number was found in HeLa cells following treatment with the electrogenic ionophore agents CCCP and valinomycin (44). Recently, acute CCCP treatment was found to promote lysosomal biogenesis through activation of transcription factor EB (TFEB) (23, 45). TFEB regulates both mitochondrial and lysosomal biogenesis and is translocated to the nucleus upon starvation, mitophagy induction and when lysosomal function is impaired (46–48). Whether TFEB translocation/activation is responsible for the increased lysosomal content in *PARK2* KO neurons remains to be examined.

Another contributing factor to the increased lysosomal content in *PARK2* KO neurons could be an impaired lysosomal degradation capacity and the build-up of undegraded cargo, as reported in iPSC-derived dopaminergic neurons from GBA patients (47, 49). Applying TEM we detected an increased number of large electron-lucent lysosomes in *PARK2* KO neurons. These less electron-dense lysosomes might represent lysosomes in which digestion of the contents is perturbed, as the normally dark lumen is a result of enzymes actively degrading the material in acidic lysosomal interior (31). Similar observations of enlarged and electron-lucent lysosomal vacuoles are described in lysosomal storage disorders (50–52). Supporting this notion, the overall lysosomal enzyme activity was significantly decreased in the *PARK2* KO neurons. A recent study on cellular models of mitochondrial dysfunction induced by chemical stressors or deletion of mitochondrial proteins including PINK1, also documented impaired lysosomal enzyme activity. These changes were dependent on ROS levels, as antioxidant treatment could revert the observed phenotypes (27). Given that the *PARK2* KO neurons have decreased levels of antioxidant defense enzymes (29), a similar mechanism could be relevant in the observations made in the present study.

The impaired lysosomal enzyme activity found here is a likely cause of the compromised autophagic flux in the *PARK2* KO neurons. However, as indicated by the proteomic data perturbations in levels of proteins involved in the autophagic process including LC3, which was significantly decreased, were also present. Decreased autophagic flux has similarly been documented in a recent study performed on *PARK2*-PD fibroblasts as well as in iPSC-derived neurons from GBA- and sporadic PD patients (47, 53, 54).

Interestingly, the GCase activity was significantly increased in the *PARK2* KO neurons whereas activity of β-Gal and overall lysosome enzymatic activity were reduced. This points to the activation of a compensatory mechanisms to increase GCase activity, as the total protein level of GCase was unchanged. A direct interaction between parkin and GCase could explain such link, as parkin can directly interact with and mediate degradation of mutant GCase (55). However, this was not the case for wildtype GCase and further studies are needed in order to address the interplay between GCase activity and parkin disfunction.

In conclusion, our results indicate that the loss of *PARK2* causes several lysosomal perturbations affecting their abundance, morphology, content and activity. Taken together with our previous data, these findings contribute to causally link two major pathological features of PD, namely mitochondrial defects and lysosomal dysregulation, supporting the idea of a direct pathogenic positive feedback between the two organelles. Future efforts must be focused on the identification of the molecular mechanistic pathways connecting mitochondrial and lysosomal perturbations with the aim of developing therapeutic approaches for PD.

## Acknowledgements

This work was supported by the Innovation Fund Denmark (BrainStem), H. Lundbeck A/S, the Danish Parkinson Foundation, the A.P. Møller Foundation for the Advancement of Medical Science, and the Faculty of Health Sciences at University of Southern Denmark. Work at the A.M.S. group at the CBMSO-Madrid was supported by grants from MINECO SAF-2017-83241-R, and ISC-III RETICS TerCel RD16/0011/0032.

The live imaging experiments reported in this paper were performed at DaMBIC, a bioimaging research core facility at the University of Southern Denmark. DaMBIC was established by an equipment grant from the Danish Agency for Science Technology and Innovation and by internal funding from the University of Southern Denmark. (+ Clair Gudex)

The authors would like to thank Dorte Lyholmer, Nadine Nadine Becker-von Buch, Ulla Melchior Hansen and Maria Pihl for excellent technical assistance.

## Materials and Methods

### CCCP (*cyanide m-chlorophenylhydrazone*) and BAF (bafilomycinA1) treatments

20 mM stock solution of CCCP (Sigma-Aldrich) was prepared by dissolving in DMSO (Sigma-Aldrich) and further diluted to get a final concentration of 10 μM. Cells were treated with CCCP or with equivalent dose of vehicle (DMSO) for 48h. CCCP solution was prepared on the day of the experiment and was protected from light.

Bafilomycin A1 (Sigma-Aldrich) was dissolved in DMSO (Sigma-Aldrich). Cells were treated with 10 nM bafilomycin A1 or with equivalent dose of vehicle (DMSO) for 6h.

### *In vitro* propagation and differentiation of neural stem cells (NSCs)

*PARK2* KO and healthy isogenic control NSC cell lines were provided by XCell Science Inc. (CA, USA). NSC lines were propagated according to well-established and standard protocol, using Geltrex-(Thermo Fisher) coated plates in Neurobasal Medium (Thermo Fisher) supplemented with NEAA, GlutaMax-I, B27, supplement (Thermo Fisher), penicilin-streptomycin, and bFGF. Cells were enzymatically passaged with Accutase (Thermo Fisher) when reaching 80-90% confluency. NSCs were differentiated according to a commercially available dopaminergic differentiation kit (#DD-001) from XCell Science Inc. (CA, USA). Differentiation was divided in two parts: an induction phase, where NSCs were differentiated into dopaminergic precursors and a maturation phase, where the dopaminergic precursor cells were differentiated into mature dopaminergic neurons. The differentiations were carried out at 37°C in a low O_2_ environment (5% CO_2_, 92% N_2_ and 3% O_2_). The cells were seeded onto poly-L-ornithine (Sigma) and laminin- (Thermo Fisher)coated wells at the density of 50.000 cells/cm^2^. Complete DOPA Induction Medium (XCell Science) supplemented with 200 ng/ml human recombinant Sonic Hedgehog (Peprotech) was changed every second day for the first 9 days of differentiation. The cells were passaged at day 5 and 10 and seeded at a desired cell density. The medium was switched to Complete DOPA Maturation Medium A and B (XCell Science) at day 10 and 16, respectively.

### Quantitative Polymerase Chain Reaction (qPCR)

Cell lysates were analyzed by qPCR for the mRNA expression of the dopaminergic markers EN1, NURR1 and GIRK2. Medium was aspirated from the wells and 500 μl cold Trizol lysis reagent (Life Technologies) was added per well. Cells were harvested using a sterile cell scraper and transferred to RNase-free eppendorf tubes. Samples were vortexed for 30 min to completely lyse the cells. RNA purification was performed through columns according to manufacturer’s instructions (ENZA total RNA Kit I, VWR). Reverse transcription from mRNA to cDNA was performed using the Super Script III reverse transcriptase kit (Invitrogen) according to manufacturer’s instruction. cDNA was processed by qPCR analysis using a protocol customized according to the instructions of the SsoFastTM EvaGreen® Supermix (BioRad) in the BioRad CFX-96 equipment. The efficiency of all primer sets was tested prior to qPCR analysis. For that, cDNA from a positive control (iPSC-derived midbrain dopaminergic neurons, day 30 of differentiation) was tested in 4 different concentrations: 1:1, 1:2, 1:5 and 1:10 and the efficiency assessed using the BioRad software. Acceptable range was defined as 100% ± 10. For specificity testing, the positive control and a negative iPSC control were included in the analysis. The expression level of the dopaminergic markers was quantified relative to the expression of the three housekeeping genes GAPDH, 18S, HPRT. Relative gene expression was assessed using the following TagMan assays: *GAPDH*, Hs02758991_g1; *18S*, Hs03003631_g1; *HPRT*, Hs02800695_m1; *EN1*, Hs00154977_m1; *NURR1*, Hs00428691_m1; *GIRK2*, Hs00158423_m1.

### Cell collection for mass spectrometry-based proteomic

*PARK2* KO and control neuronal cultures from 3 independent differentiations were collected on ice in phosphate-buffered saline (PBS, ThermoFisher) with protease-(Complete tablets, Roche) and phosphatase inhibitors (PhosSTOP tablets, Roche). The QProteome Mitochondria Isolation Kit (Qiagen #37612) was used according to the manufacturer’s instruction for standard preparation.

### Lysis, reduction and enzymatic digestion

Samples were sonicated for two times ten seconds at 50% amplitude on ice and incubated for 30 min at room temperature (RT) in lysis buffer consisting of 6 M urea (Sigma), 2 M thiourea (Sigma), 20 mg/ml sodium dodecyl sulfate (SDS, GE Healthcare), 40 nM N-ethylmaleimide (NEM, Sigma) and protease inhibitor. The proteins were denatured and reduced in 6 M urea, 2 M thiourea and 10 mM TCEP (Thermo Fisher) at RT. After vortexing, the samples were incubated at RT for 2 hours with 1 μl endoproteinase Lys-C (Wako). The samples were diluted 10 times in 20 mM TCEP and 20 mM tetraethylammonium bromide (TEAB, Sigma), pH 7.5, and sonicated followed by digestion with 1 μg trypsin (Sigma) per 50 μg peptide over night (ON) at RT.

### Desalting, Tandem Mass Tag labelling and enrichment of phosphorylated peptides

The samples were acidified with 0.1% trifluoroacidic acid (TFA, Sigma) and desalted using two self-made P200-tip-based columns per sample. 45 μg of each sample were labeled with Tandem Mass Tag (TMT) Sixplex Isobaric label Reagents (ThermoFisher) according to the manufacturer’s instructions. Efficient labeling was confirmed by MALDI, and the labeled peptides mixed 1:1:1:1:1:1 and dried.

### Hydrophobic interaction liquid chromatography (HILIC) and high pH fractionation

Mono-phosphorylated and non-modified peptides were fractionated to reduce sample complexity using HILIC. To increase the coverage, high pH fractionation was also performed using approximately 50 μg peptide of the non-modified peptide sample. Briefly, the sample was dissolved in 1% ammonium hydroxide (NH_3_, Sigma), pH 11, and loaded on a R2/R3 column equilibrated with 0.1% NH_3_. The peptides were eluted in a stepwise fashion using a gradient of 5%-60% ACN/0.1% NH_3_. All fractions were dried by vacuum centrifugation.

### Reversed-phase nano LC-ESI-MS/MS

The samples were resuspended in 0.1% formic acid (FA) and loaded onto a two-column EASY-nLC system (Thermo Scientific). The pre-column was a 3 cm long fused silica capillary (100 μM inner diameter) with a fritted end and in-house packed with ReproSil - Pur C18 AQ 5 μm (Dr. Maisch GmbH) whereas the analytical column was a 17 cm long fused silica capillary (75 μm inner diameter) and packed with ReproSil–Pur C18 AQ 3 μm reversed-phase material (Dr. Maisch GmbH). The peptides were eluted with an organic solvent gradient from 100% phase A (0.1% FA) to 34% phase B (95% ACN, 0.1% FA) at a constant flowrate of 250 nL/min. Depending on the samples based on the HILIC, the gradient was from 1 to 30% solvent B in 60 min or 90 min, 30% to 50% solvent B in 10min, 50%-100% solvent B in 5 min and 8 min at 100% solvent B. The nLC was online connected to a QExactive HF Mass Spectrometer (Thermo Scientific) operated at positive ion mode with data-dependent acquisition. The Orbitrap acquired the full MS scan with an automatic gain control (AGC) target value of 3×10^6^ ions and a maximum fill time of 100ms. Each MS scan was acquired at high-resolution (120,000 full width half maximum (FWHM)) at m/z 200 in the Orbitrap with a mass range of 400-1400 Da. The 12 most abundant peptide ions were selected from the MS for higher energy collision-induced dissociation (HCD) fragmentation (collision energy: 34V). Fragmentation was performed at high resolution (60,000 FWHM) for a target of 1×10^5^ and a maximum injection time of 60ms using an isolation window of 1.2 m/z and a dynamic exclusion. All raw data were viewed in Thermo Xcalibur v3.0.

### Mass spectrometry data analysis

The raw data were processes using Proteome Discoverer (v2.1, ThermoFisher) and searched against the Swissprot human database using an in-house Mascot server (v2.3, Matrix Science Ltd.) and the Sequest HT search engine. Database searches were performed with the following parameters: precursor mass tolerance of 10 ppm, fragment mass tolerance of 0.02 Da (HCD fragmentation), TMT 6-plex (Lys and N-terminal) as fixed modifications and a maximum of 2 missed cleavages for trypsin. Variable modifications were NEM on Cys and N-terminal acetylation along with phosphorylation of Ser/Thr/Tyr for the phosphorylated group. Only peptides with up to a q-value of 0.01 (Percolator), Mascot rank 1 and cut-off value of Mascot score > 15 were considered for further analysis. Only proteins with more than one unique peptide were considered for further analysis in the non-modified group.

### Sample preparation for transmission electron microscopy (TEM)

The TEM sample preparation procedure included five major steps: fixation, dehydration, infiltration and embedding, sectioning, and contrasting. Cells were seeded on 13 mm Thermanox plastic coverslips (Nunc) with a density of 80.000 cells/cm^2^. At day 25 of differentiation cell were primarily fixed in 3% gluteraldehyde (Merck) in 0.1 M Natrium phosphate buffer with pH 7.2 at 4°C for 1h and stored in a 0.1 M Na-phosphate buffer at 4°C until further analysis. When ready, the cells were embedded in 4% agar at 45°C (Sigma) under the stereomicroscope and cut into 1-2 mm^3^ blocks, which were then washed with 0.1 Na-phosphate buffer, followed by a post-fixation in 1% osmium tetroxide (EMS) in 0.1 Na-phosphate buffer (pH 7.2) for 1h at RT. Cells were washed in MilliQ water, followed by a stepwise dehydration in a series of ascending ethanol concentrations ranging from 50% EtOH to 99% EtOH). Then propylene oxide (Merck) was used as an intermediate allowing for infiltration with Epon (812 Resin, TAAB). The following day, the agar blocks were placed in flat molds in pure Epon, which was cured at 60°C for 24h. Approximately 8 semi-thin sections (2 μm) from one block were cut on an ultramicrotome with glass knife (Leica, Reichert Ultracut UTC). These were stained with 1% toluidine blue in 1% Borax and evaluated by light microscopy to locate areas with adequate number of cells for further processing. Ultra-thin sections (70 nm) were cut on the ultramicrotome with a diamond knife (Jumdi, 2 mm), and the ultra-thin sections were collected onto TEM copper grids (Gilder), and then stained with 2% uranyl acetate (Polyscience) and lead citrate (Reynolds 1963). The samples were then evaluated and the image database collected using a Philips CM100 transmission electron microscope equipped with a Morada digital camera equipment and iTEM software system.

### Measurements of lysosomal numbers on TEM images

Six TEM grids from each line were used for analysis, and at low magnification 10 spots were randomly chosen. Images of each spot were captured using high magnification (19.000X). To estimate lysosomal number, all lysosomes were counted on each micrograph. For further morphometric analysis individual organelles were selected and processed manually using the different selection tools in ImageJ software.

### Western blotting

Cell lysates were analyzed by sodium dodecyl sulphate polyacrylamide gel electrophoresis (SDS-PAGE) to separate the proteins by molecular weight followed by western blotting. Cells were differentiated in 6-well plates (Costar) with a cell density of 50.000 cells/cm^2^ and harvested using accutase. Cell pellets were resuspended in PBS containing phosphatase (PhosphoSTOP tablet, Roche) and protease inhibitors (Complete Mini Tablets, Roche) on ice. Sample were centrifuged (10.000 rpm, 5 min, 4°C), supernatant aspirated and cells sonicated for 2×10 sec at amplitude 2, low speed on ice in PBS containing phosphatase and protease inhibitors supplemented with 1% Triton X-100. Afterwards, samples were incubation for 30 min at 4°C with shaking and then centrifuged (10.000 rpm, 15 min, 4°C) and the supernatants were collected. The protein content in the samples was determined using the Micro Pierce® Bicinchoninic acid (BCA) Protein Assay (Thermo Scientific). Prior to electrophoresis, equal amounts of protein from each sample were mixed 1:1 with NuPAGE reducing loading buffer and denatured for 10 min at 70°C. For detection of TH 20 μl (5 μg protein)/well sample and 10 μl/well SeeBlue Plus2® prestained standard (Novex) were loaded on 4-12% Bis/Tris gels (Invitrogen NuPAGE). Electrophoresis was performed with NuPAGE MOPS SDS running buffer (Invitrogen NuPAGE) supplemented with 0.25% antioxidant (Invitrogen NuPAGE) for 50 min at 200 V. For detection of MAP2 20 μl (5 μg protein)/well sample and 10 μl/well HiMark prestained standard (Novex) were loaded on 3-8% Tris-Acetate gels (Invitrogen NuPAGE). Electrophoresis was performed with Tris-Acetate SDS running buffer (Invitrogen NuPAGE) supplemented with 0.25% antioxidant for 60 min at 150 V. Proteins were blotted from the gels to PVDF membranes (Invitrogen) using the IBlotTM Dry blotting system from Invitrogen. Blotting was performed for 7 min for TH and 9 min for MAP2. Membranes were blocked for 60 min in 5% milk powder and 0.05% Tween-20 in 0.05M TBS at 4°C. After a short wash in 0.05% Tween-20 in 0.05M TBS, the membranes were incubated ON with primary antibodies and the following dilutions were used: mouse anti-TH (#MAB5280 Chemicon/Millipore) 1:2000, mouse anti-MAP2a+b (#M1406, Sigma) 1:500, rabbit anti-LAMP1 (Abcam #24170) 1:500, rabbit anti-LAMP2a (Abcam #18528) 1:500, rabbit anti-GBA (Abcam #154856) 1:1000, and rabbit anti-LC3 (Cell Signaling #4599) 1:1000. After incubation with primary antibodies, the blots were washed 4 times in 0.05% Tween-20 in 0.05M TBS for 5 min at RT to remove unbound primary antibodies. Subsequently, blots were incubated with an appropriate horseradish peroxidase (HRP)-conjugated secondary antibody (#P0260 or #P0217, DAKO) with dilution 1:2000 for 1h at RT. Unbound secondary antibodies were afterwards removed by washing the blots 4 times in 0.05% Tween-20 in 0.05M TBS for 5 min at RT, followed by chemiluminescence development (ECL kit, ThermoFisher Scientific) and visualization using the ChemiDoc MP imaging system (BioRad). As loading control, all blots were subsequently incubated with α-actin antibody (mouse anti-α-actin, #MAB1501, Chemicon/Millipore; 1:6000) ON at 4°C and developed as described above. Densitometric analyses of band intensities were quantified using Image Lab software (BioRad) and protein expression levels were normalized to α-actin.

### Immunofluorescence staining

Cells cultured on coverslips in 24-well plates (Costar) were fixed in 4% paraformaldehyde (PFA, Sigma) in 0.15 M Sørensen buffer (potassium dihydrogen phosphate (Merck) and disodium phosphate (Merck)) for 20 min at RT and then washed 2 times in 0.15 M Sørensen Buffer for 15 min. Fixed cells were washed once in 0.05 M Tris Buffered Saline (TBS, Sigma) for 5 min at RT and then permeabilized by washing 3 times in 0.1% Triton X-100 (Sigma) in 0.05 M TBS for 15 min at RT, followed by blocking in 5% goat serum (Millipore) in 0.05 M TBS for 30 min at RT to avoid non-specific binding of the primary antibodies. Afterwards, primary antibodies diluted in 5% goat serum were added to the cells and incubated ON at 4°C. Primary antibodies were used in the following concentrations: rabbit anti-tyrosine hydroxylase (TH, Millipore #AB152) 1:600, mouse anti-microtubule-associated protein 2a+b (MAP2, Sigma #M1406) 1:2000, rabbit anti-LAMP1 (Abcam #108597) 1:1000. After ON incubation with primary antibody, cells were washed 3 times in 0.1% Triton X-100 in 0.05 M TBS for 15 min at RT to remove unbound primary antibodies. The cells were then incubated with fluorophore-conjugated secondary antibodies: Alexa Fluor 488 goat anti-rabbit IgG (Invitrogen #A11008) or Alexa Fluor 555 goat anti-rabbit IgG (Abcam #150078) 1:500 diluted in 5% goat serum in 0.05 M TBS for 2h at RT. Unbound secondary antibodies were afterwards removed by washing the cells 2 times in 0.05 M TBS for 15 min at RT. Cells were then counterstained with 10 μM Hoechst (Sigma) for 15 min at RT to stain all nuclei. Finally, cells were mounted onto glass slides using ProLong® Diamond mounting medium (Molecular Probes).

### Image analysis

Fluorescence images were acquired on a FluoView FV1000MPE – Multiphoton Laser Confocal Microscope (Olympus) 20X or 60X magnification, in a blinded manner on 5 randomly chosen confocal fields per coverslip from independent experiments. TH+ dopaminergic neurons and MAP2+ mature neurons stainings were quantified manually in ImageJ using the Cell Counter plugin and normalized to total cell numbers as quantified by DAPI+ nuclei analysis in CellProfiler software. Only cells displaying an extensive immunostaining with a well-preserved cellular morphology were counted. Analysis lysosomal numbers was performed automatically in ImageJ software by converting LAMP1 images to binary format and analyzing particles. Total area of lysosomes was normalized to total cell numbers as quantified by DAPI+ nuclei analysis in ImageJ software.

### GCase activity assay

Cell pellets were sonicated at 10 amp for 10 sec in citrate phosphate buffer pH 5.4 consisting of 0.1 M citric acid (Sigma) and 0.2 M dibasic sodium phosphate (Sigma) with 0.25% (v/v) Triton X and 0.25% (w/v) taurocholic acid (Sigma). Samples were centrifuged at 800 g for 5 min at 4°C and supernatant collected. Following protein determination equal amounts of protein from each sample were diluted in citrate phosphate buffer in quadruplicates. One replicate of each sample was treated with 1 mM conduritol B epoxide (CBE, Calbiochem) for 10 min before all samples were incubated for 1h with 2.5 mM of the fluorescent GCase substrate methylumbillifery β-D-glucopyranoside (4MUG, Sigma) at 3°C in the dark. The reaction was quenched with 1 M glycine buffer (Sigma) pH 10.8 and the fluorescent levels analyzed on the PHERAStar FSX plate reader (BMG Labtech). Values from CBE-treated wells were subtracted as background.

### Total Lysosomal Intracellular Activity Assay

Measurement of the total lysosomal intracellular activity was performed using Lysosomal Intracellular Activity Assay Kit (Cell-Based) (BioVision) according to manufacturer’s protocol. Briefly, cells were plated on 13 mm glass coverslips in a 24-well culture plate (40.000 cells/cm^2^). Prior to the analysis the cells were incubated for 8h in medium supplemented with 10% FBS at 37°C. Then the media was removed and replaced with fresh aliquots supplemented with 0.5% FBS. 15 μl of Self-Quenched Substrate was added per 1 ml of media into each well and the plate incubated for 1h at 37°C. 10 min before the end of incubation 1:1000 Hoechst solution (Sigma) was added to each well to stain nuclei. After incubation the cells were analyzed under fluorescence microscope with 488 nm excitation filter to visualize the fluorescence of released Self-Quenched Substrate, which proportional to the total lysosomal intracellular activity. Fluorescence images were acquired on a FluoView FV1000MPE – Multiphoton Laser Confocal Microscope (Olympus) 60X magnification, in a blinded manner on 5 randomly chosen confocal fields per coverslip from independent experiments. Analysis was performed automatically in ImageJ software by converting images to binary format and analyzing particles. Total area of the staining was normalized to total cell numbers as quantified by Hoechst+ nuclei analysis in ImageJ software.

### Beta Galactosidase (β-Gal) Assay

Measurement of β-Gal activity was performed using Fluorometric Beta Galactosidase Activity Assay Kit (BioVision) according to manufacturer’s protocol. Briefly, cells were plated in a 6-well plate (60.000 cells/cm^2^) and homogenized with on ice with 100 μl ice cold β-Gal Assay Buffer and kept on ice for 10 min. Cell lysates were transferred to eppendorf tubes and centrifuged at 10.000 g, 4°C for 5 min, and subsequently supernatant was collected to a new eppendorf tubes. 30 μl of supernatant in triplicates was transferred into desired wells in a 96-well black plate. For Positive Control the β-Gal Positive Control was diluted 1:25 in the Assay Buffer and mixed well. 15 μl in triplicates of diluted β-Gal Positive Control was added into desired wells. The volume of samples and Positive Control wells was adjusted to 50 μl/well with β-Gal Assay Buffer. Required amount of Reaction Mix was prepared, containing β-Gal Assay Buffer and β-Gal Substrate and added to each well in volume of 50 μl. Fluorescence signal (Ex/Em = 480/520 nm) was measured in kinetic mode and the Fluorescein Standard Curve was read in Endpoint mode.

### Autophagic Flux

To examine the autophagic flux, we determined if LC3 is degraded in a lysosomal-dependent manner by using bafilomycin A1, an inhibitor of lysosomal acidification but also, independently of its effect on lysosomal pH, of fusion between autophagosomes and lysosomes. To this purpose, differentiated neurons were incubated with DMSO (untreated) or with medium containing bafilomycin A1 (10 nM for 6 h). Western blot analysis was performed on cell lysates using antibodies against LC3 (Cell Signaling) and α-actin (Abcam). The ratio between LC3-II and LC3-I was calculated and normalized to α-actin, through densitometric analysis using ImageJ software, and the autophagic flux was calculated as the ratio of LC3-II of cells treated with bafilomycin A1 and untreated cells.

### Statistical analysis

Statistical analysis was performed in GraphPad Prism 7.0 software using two-tailed unpaired Student’s t-test and one-way ANOVA with Tukey’s Multiple Comparison Test where appropriate. Data were considered statistically significant at p < 0.05 (*), p < 0.01 (**) and p < 0.001 (***). Data are presented as mean ± standard error of the mean (SEM).

